# Systematic Analysis of Lysine Succinylation in Vero cells infected with Small Ruminant Morbillivirus

**DOI:** 10.1101/2021.05.17.444595

**Authors:** Xuelian Meng, Xueliang Zhu, Rui Zhang, Zhidong Zhang

## Abstract

Small ruminant morbillivirus (SRMV), formerly named Peste-des-petits ruminants virus (PPRV), belongs to the genus *Morbillivirus* in the family *Paramyxoviridae* and causes a highly contagious disease in small ruminants, especially goats and sheep. Lysine succinylation is a newly identified and conserved modification and plays an important role in host cell response to pathogen infection. Here, we present a global analysis of the succinylome of SRMV-infected Vero cell using succinylation specific antibody-based enrichment and dimethylation labeling-based quantitative proteomics. As a result, 2633 succinylation sites derived from 823 cellular proteins were quantified. Comparative analysis revealed that 228 down-regulated succinylation sites on 139 proteins and 44 up-regulated succinylation sites on 38 proteins were significantly modified in response to SRMV infection, seven lysine succinylation motifs were identified. Bioinformatics analysis showed that the succinylated proteins mainly participated in cellular respiration and biosynthetic process. Protein-protein interaction networks of the identified proteins provided further evidence that a variety of ATP synthase subunits and carbon metabolism were modulated by succinylation while the overlapped proteins between succinylation and acetylation are involved in glyoxylate and dicarboxylate metabolism. Taken together, these findings provide the first report of succinylome in SRMV infection, lysine acetylation may have a more important effect than succinylation in SRMV infection. These findings provide a novel view on investigating the pathogenic mechanism of SRMV.

## INTRODUCTION

Peste-des-petits-ruminants virus (PPRV) is the etiological agent of Peste des petits ruminants (PPR), which is a highly contagious transboundary disease and included in the OIE (the World Organization of Animal Health) list of notifiable terrestrial animal diseases (1, 2). In 2016, according to the latest virus taxonomy of the International Committee on Taxonomy of Viruses, PPRV was renamed small ruminant morbillivirus (SRMV) which belongs to the *Morbillivirus* genus in family *Paramyxoviridae*, alongside measles morbillivirus (MV), rinderpest morbillivirus (RPV), canine morbillivirus (CDV), phocine morbillivirus (PDV), cetacean morbillivirus (CeMV) and feline morbillivirus (FeMV) (3–5). SRMV has a negative, non-segmented single-stranded genomic RNA encoding six structural proteins, nucleocapsid protein (N), phosphoprotein (P), matrix protein (M), fusion protein (F), haemagglutinin protein (H) and large polymerase protein (L) and two non-structural proteins (V and C protein). SRMV was first reported in the Ivory Coast in 1942 and is now present in over 70 countries across Asia, Africa, and the Near and Middle East, and has also been detected in Europe (Bulgaria) (6), inhibiting trade and causing significant economic losses. Following the successful eradication of rinderpest, the OIE and FAO have proposed a strategy for the control and eradication of PPR by 2030 (7).

Host proteins are highly contested targets in the ongoing ‘arms race’ between viruses and the host. Viruses “hijack” host cellular functions to facilitate their replication and inhibit host antiviral defenses. On the contrary, the host actively mobilizes the immune antiviral response to resist viral invasion, or inhibit the replication of viral particles, or eliminate virus particles. In previous studies, host cell responses to infection with various viruses were deciphered by clustered regularly interspaced short palindromic repeat (CRISPR) (8–10), small interfering RNA (siRNA)(11) and transcriptomic and proteomic analyses (12, 13). Today, it is well known that protein post-translational modifications (PTMs) significantly affect the diverse function of proteins through the modulation of biological processes, protein activity, cellular location and protein-protein interaction (PPI) by transfer modified groups to one or more amino acid residues. In recent years, protein PTMs have become a hot topic in viral infection.

To date, over 450 protein modifications including more than 200 PTMs have been identified (14). They are dynamic and reversible protein processing events that play key roles in the response to the pathogenesis and development of diseases (15–18). Some PTMs including phosphorylation, methylation, acetylation and succinylation have been shown to potently regulate innate immunity and inflammation in response to virus infectious (19, 20). Of the 20 amino acid residues, lysine is a frequent target of covalent modifications because it can accept different types of chemical groups. Lysine succinylation is a newly discovered and meagerly studied modification that is evolutionarily conserved from bacteria to mammals. It can transfer a larger structural moiety (a succinyl group, -CO-CH2-CH2-CO-) than that in acetylation, and changes the charge on the modified residues from +1 to −1 causing a two-unit charge shift in the modified residues. Consequently, the structure and function of succinylated protein might have greater changes, which in turn results in substantial changes in the chemical properties of the target proteins indicating that succinylation can potentially regulate the protein structure and function associated with diverse cellular processes such as metabolism and translation (21–23).

Lysine succinylation was first reported in the active site of homoserine transsuccinylase (24). So far, lysine succinylation is widespread in diverse species ranging from prokaryotes and eukaryotes, from plant to animal (14, 25–28). It causes unique functional consequences and affects chromatin structure, conformational stability and gene expression (23, 29). In recent years, more and more attention has been paid to the high coincidence of lysine succinylation modification with acetylation modification (28, 30–35). The majority of succinylation sites in bacteria, yeast and mammal cell were acetylated at the same position, which suggest that these two modifications may affect the performance of proteins together. The coincidence sites of lysine succinylation were also mostly located in polar, acid or alkaline amino acid regions, and were exposed to protein surface (23, 24, 28). Moreover, succinylation occurs at a low level and as such, many succinylation sites remain unidentified. Since the fundamental mechanisms for succinylation are still unknown, more efforts are necessary to find the mechanism.

Despite the advanced findings at a quick pace with the developments in mass spectrometry (MS) technologies and peptide enrichment methods, global succinylome and the relationship between succinylation and acetylation remain an understudied aspect of SRMV infection. The contribution of succinylation to host defense or virus replication is still unknown. Therefore, systematical analyses of host cellular protein succinylation might provide helpful insights into SRMV pathogenesis and replication that could lead to the discovery of antiviral targets.

In the present study, we aimed to systematically investigate the succinylome of Vero cells with or without SRMV infection, by combining dimethylation labeling and LC-MS/MS analysis. we successfully quantified 2633 succinylation sites on 823 proteins with diverse molecular functions, biological processes and subcellular localizations. 272 succinylation sites on 177 proteins were significant modified in response to SRMV infection, seven lysine succinylation motifs were identified. PPI networks further indicated that a variety of ATP synthase subunits and carbon metabolism were modulated by succinylation, the overlapped proteins between succinylation and acetylation were involved in glyoxylate and dicarboxylate metabolism. Overall, the results provided the first extensive dataset of lysine succinylation in Vero cells infected by SRMV, and a novel view on investigating the infection mechanism of SRMV.

## RESULTS

### Global detection of lysine succinylated sites on SRMV-infected Vero cellular proteins

To explore and identify the host proteins with acetylated sites involved in the SRMV replication process, we detected the abundance of acetylated proteins in SRMV-infected Vero cells by western blotting and chose 24 hpi as the time point for quantitative proteomic analysis (Fig. 1). Repeatability among the 3 biological replicates ranged from 0.41 to 0.50 for the succinyl-proteome. The near-zero distribution of mass error and that the errors were predominantly < 0.02 Da (Fig. 2). Most of the enriched lysine-succinylated peptide lengths were in the range of 7-21 segments, which are consistent with cutting by trypsin at lysine residue sites.

**Fig.1.**
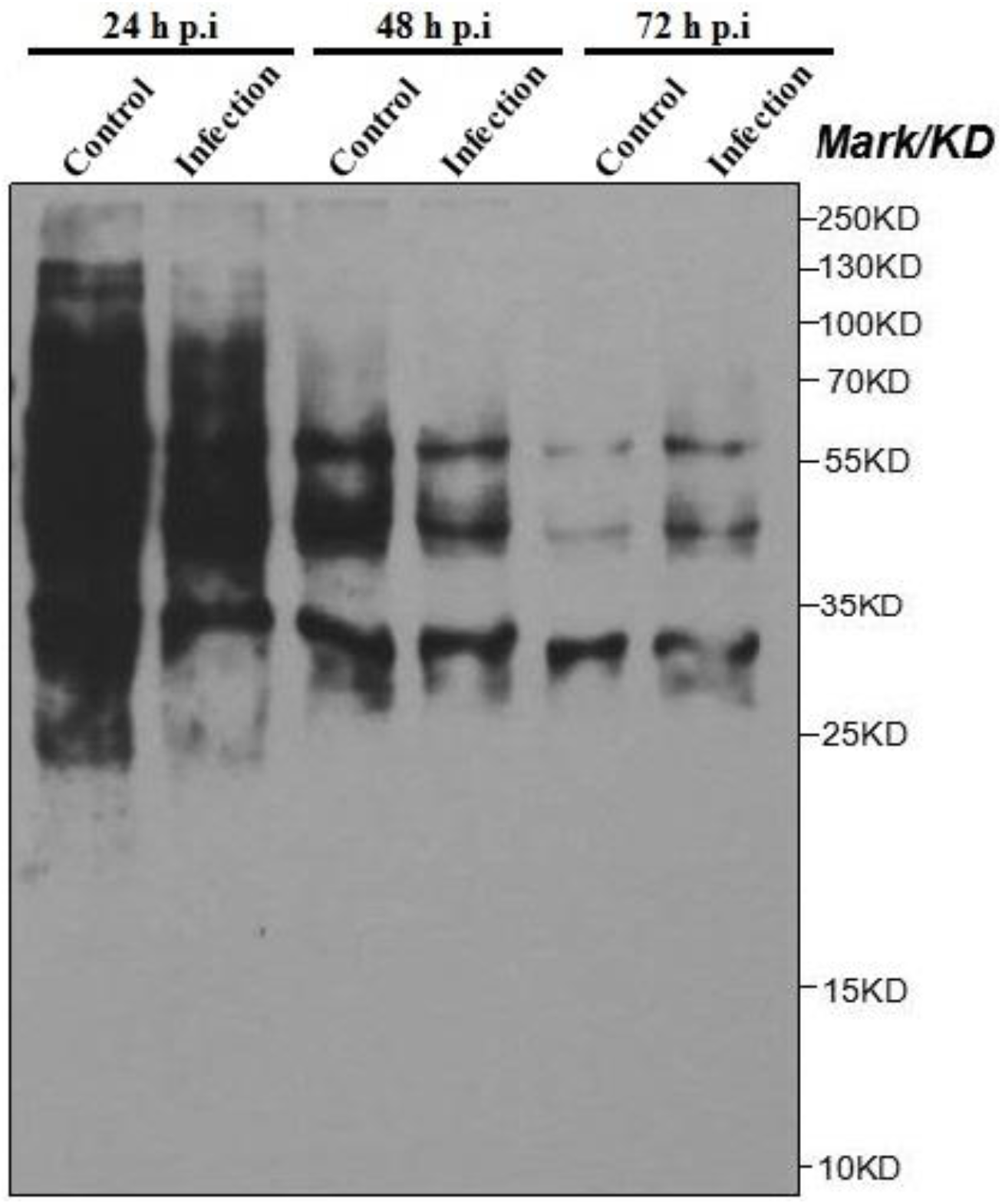
Western blotting with anti-pan succinyllysine antibody in response to SRMV infection in Vero cell

**Fig.2.**
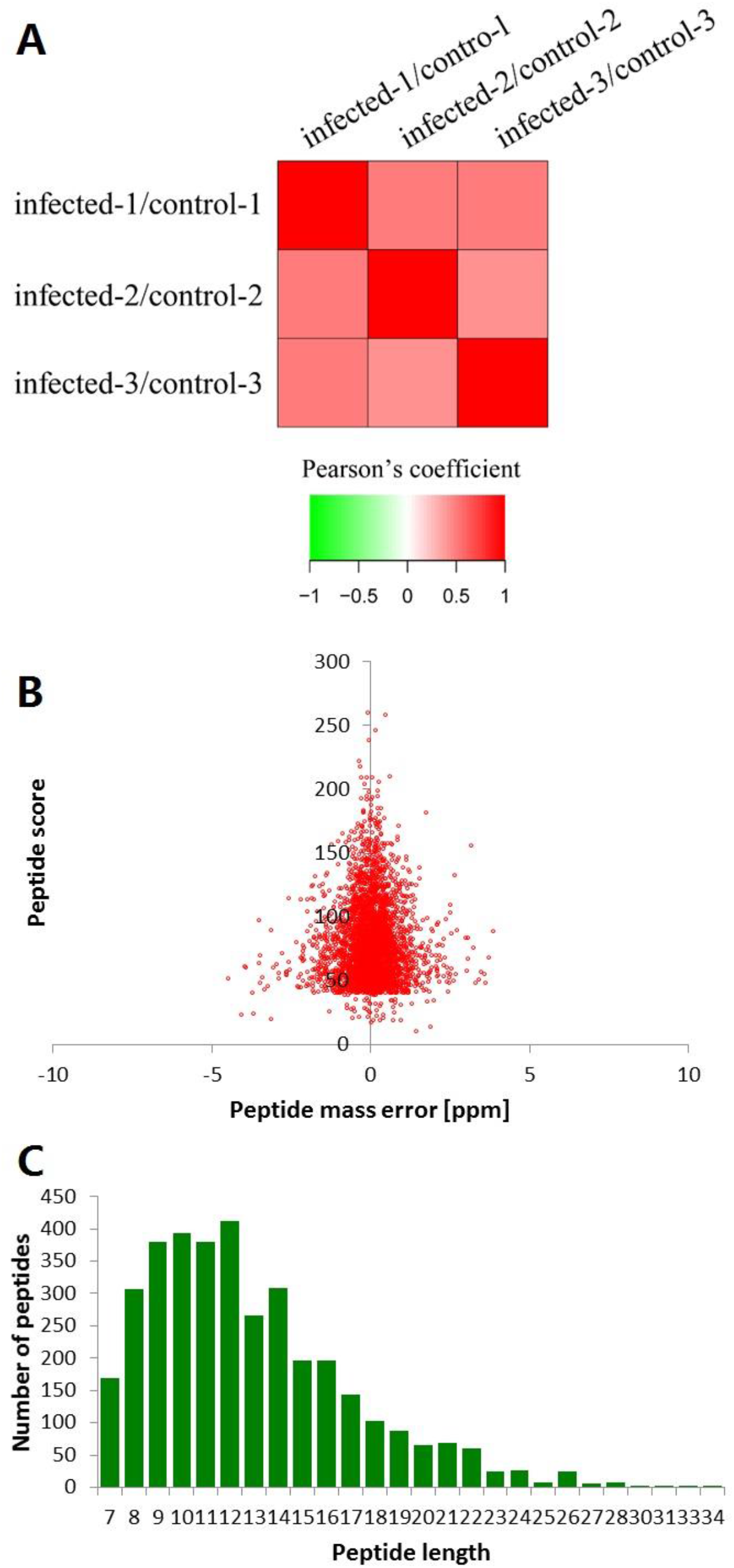
Pearson’s correlation analysis for succinyl-proteome and QC validation of MS data A. Pearson’s correlation of protein quantitation; B. Mass delta of all identified peptides; C. Length distribution of all identified peptides

A total of 2840 lysine-succinylated sites across 875 proteins were identified, of which 2633 sites on 823 proteins were quantified (Supplementary Table S1). The number of succinylated sites per protein varies from 1 to 33 depending on the protein. Among these proteins, about 383 (43.77%) included a single succintylation site, 176 (20.11%) included two sites, 86 (9.83%) included three sites, 62 (7.09%) included four sites, and 168 (19.20%) included five or more than five sites (Fig. 3). 28 lysine succinylation sites were found on histone proteins, including 1 site on H1C, 2 sites on H2A, 16 sites on H2B, 4 sites on H3 and 5 sites on H4.

**Fig.3.**
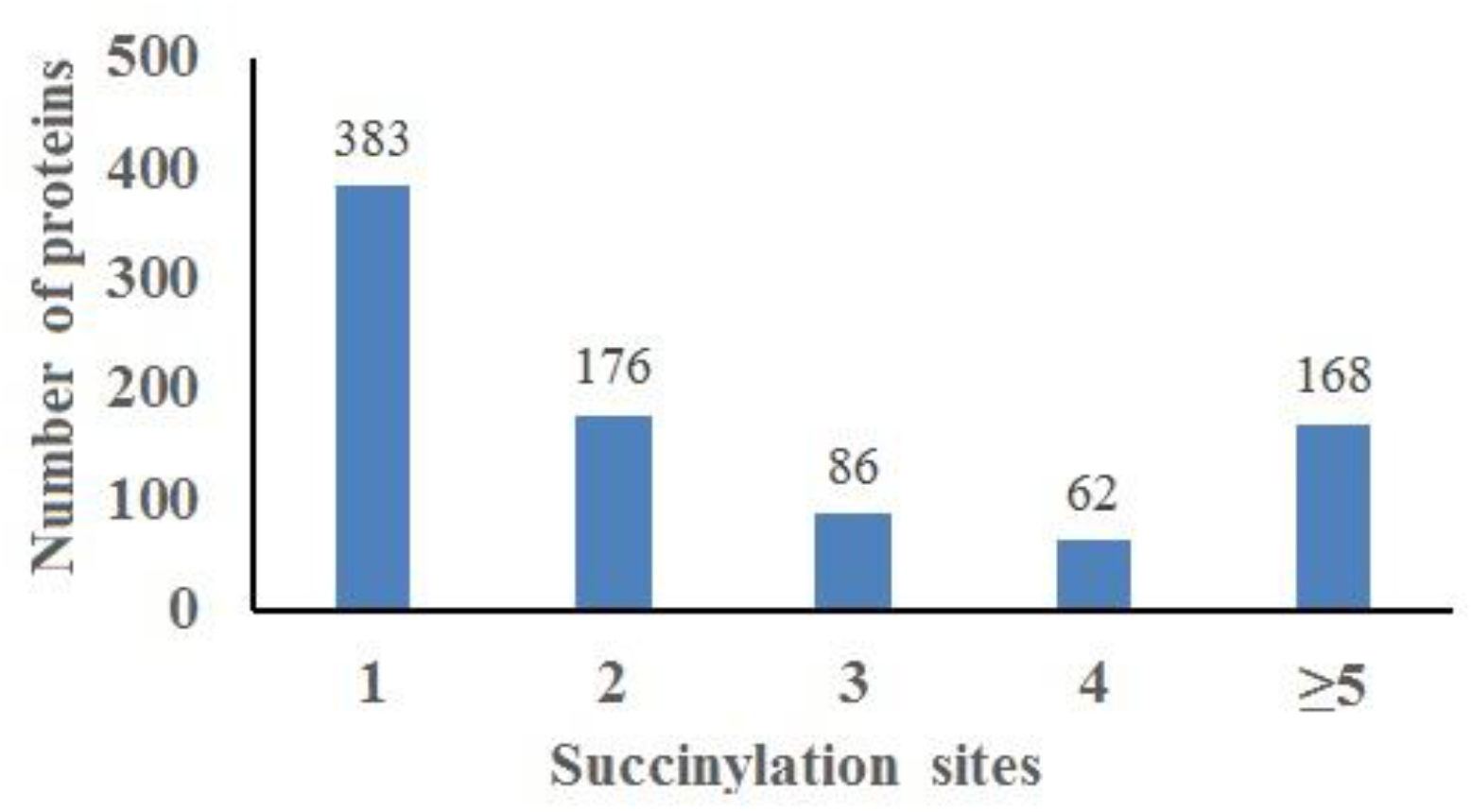
Profile of the identified succinylated sites and proteins in SRMV-infected vero cell

Based on a threshold of 1.2 fold changes and t test p< 0.05 as standards, among the 2633 quantifiable sites, 272 succintylated sites on 177 proteins can response to SRMV infection (Fig. 4, Supplementary Table S2). 44 succinylated sites on 38 proteins were quantified as significantly upregulated, and 228 succinylated sites on 129 proteins were significantly downregulated. Most of these proteins were modified at single succinylation site, including clathrin heavy chain (Lys557), CCT3 (Lys 353), CCT5 (Lys 223), ezrin (Lys 57), plectin (Lys 3390) and NLRX1 (Lys 656). 58 proteins were modified at multiple sites, including heat-shock protein (Hsp) A9 (7 sites), HspA5 (4 sites), Hsp90AB1 (3 sites), HspA8 (2 sites) and vimentin (2 sites).

**Fig.4.**
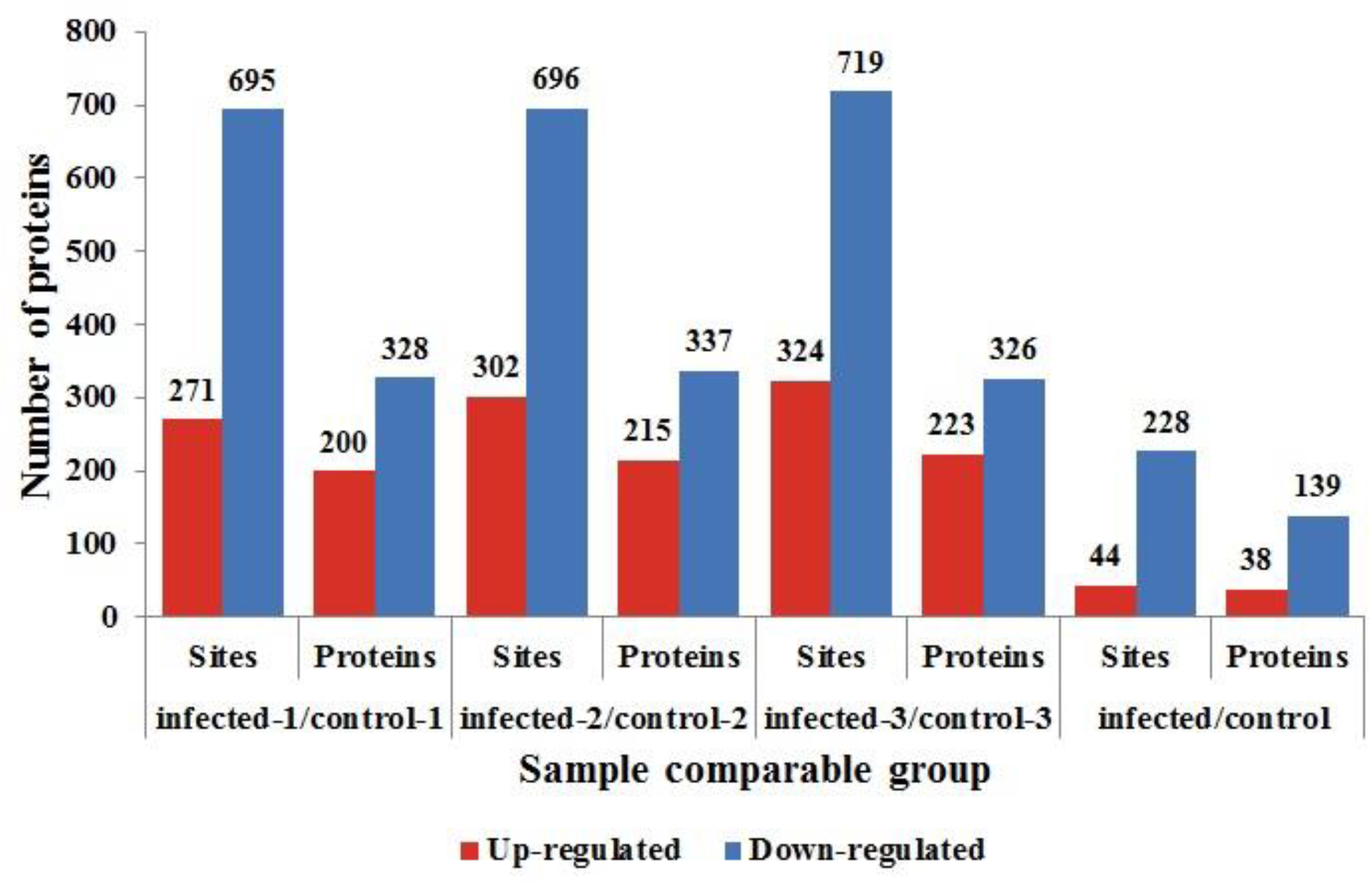
Summary of differentially quantified succinylated sites and proteins in different comparable group

### Motifs of succinylated peptides

To examine the properties of the succinylated lysine in SRMV-infected Vero cell, the sequences from the −10 to +10 positions flanking the succinylated sites were investigated using the Motif-X program with a significance threshold of p<0.000001. The amino acids flanking the succinylated sites were matched to the whole size, and the motif enrichment was illustrated in the form of a heat map.

Of all succinylated peptides, seven significantly enriched lysine succinylation site motifs from 1310 modified sites were identified (Fig. 5). These motifs are AKsc, Ksc******K, Ksc*******K, Ksc***K*****V, A****Ksc, Ksc******V and V*Ksc (Ksc: succinylated lysine; *: residue of a random amino acid). According to the position and other properties of the residues around the succinylated lysine, lysine (K) at +7/+8 or alanine (A) at −1 position was more readily succinylated, the frequency of K at +4 and valine (V) at +10 position was the lowest. The positively charged lysine (K) residue was enriched at the positions +4 to +8. These results showed that there was a preference for aliphatic amino acids around succinylation sites.

**Fig.5.**
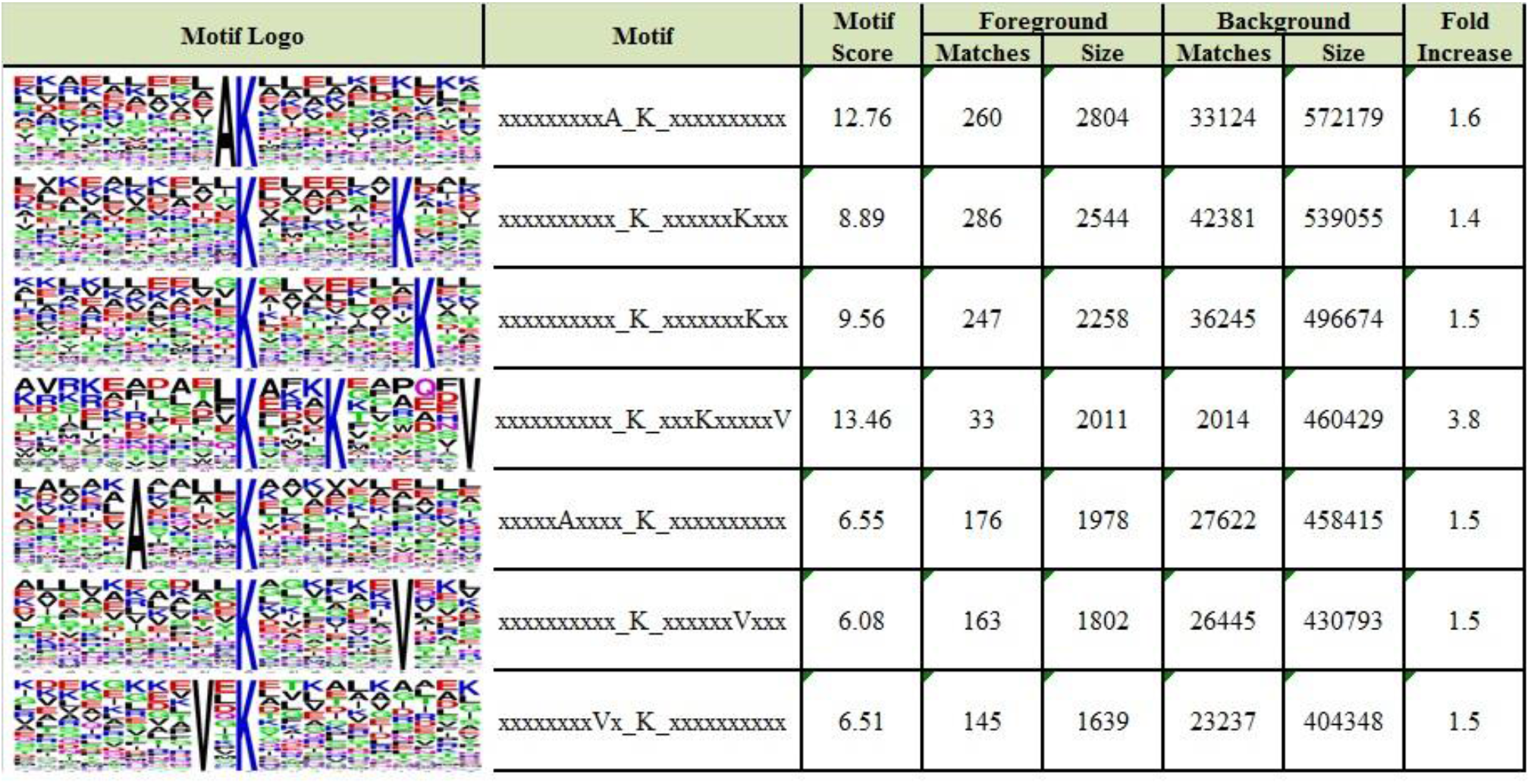
The specific motifs of succinylated peptides.

### Functional classification of the differential succinylated proteins

All of the differential succinylated proteins (DScPs) were classified by GO functional classification analysis based on their biological process, molecular function, subcellular localization and COG/KOG categories (Supplementary Table S3). In the biological processes classification, three major classes of succinyl-proteins were associated with metabolic, cellular and single-organism processes, accounting for 35%, 29% and 23% of the total DScPs, respectively (Fig. 6A). Cell (41%), organelle (22%) and membrane (20%) were identified as favorable cellular component for succinylated proteins (20%) (Fig. 6B). The succinylated proteins were mostly related to binding catalytic activity (49%) and binding (37%) (Fig. 6C). Most of DScPs were more abundant in mitochondria (54%), cytoplasm (25%), extracellular (6%) and nucleus (6%) (Fig. 6D). Moreover, the results of COG/KOG classification showed that a total of 166 DScPs were successfully annotated into 4 categories (Fig. 7). 78 DScPs (47%) were involved in metabolism, and 41% of these proteins played roles in energy production and conversion. 41 DScPs (25%) were related to cellular processes and signaling, and 71% of these proteins were involved in post-translational modification, protein turnover and chaperones.

**Fig.6.**
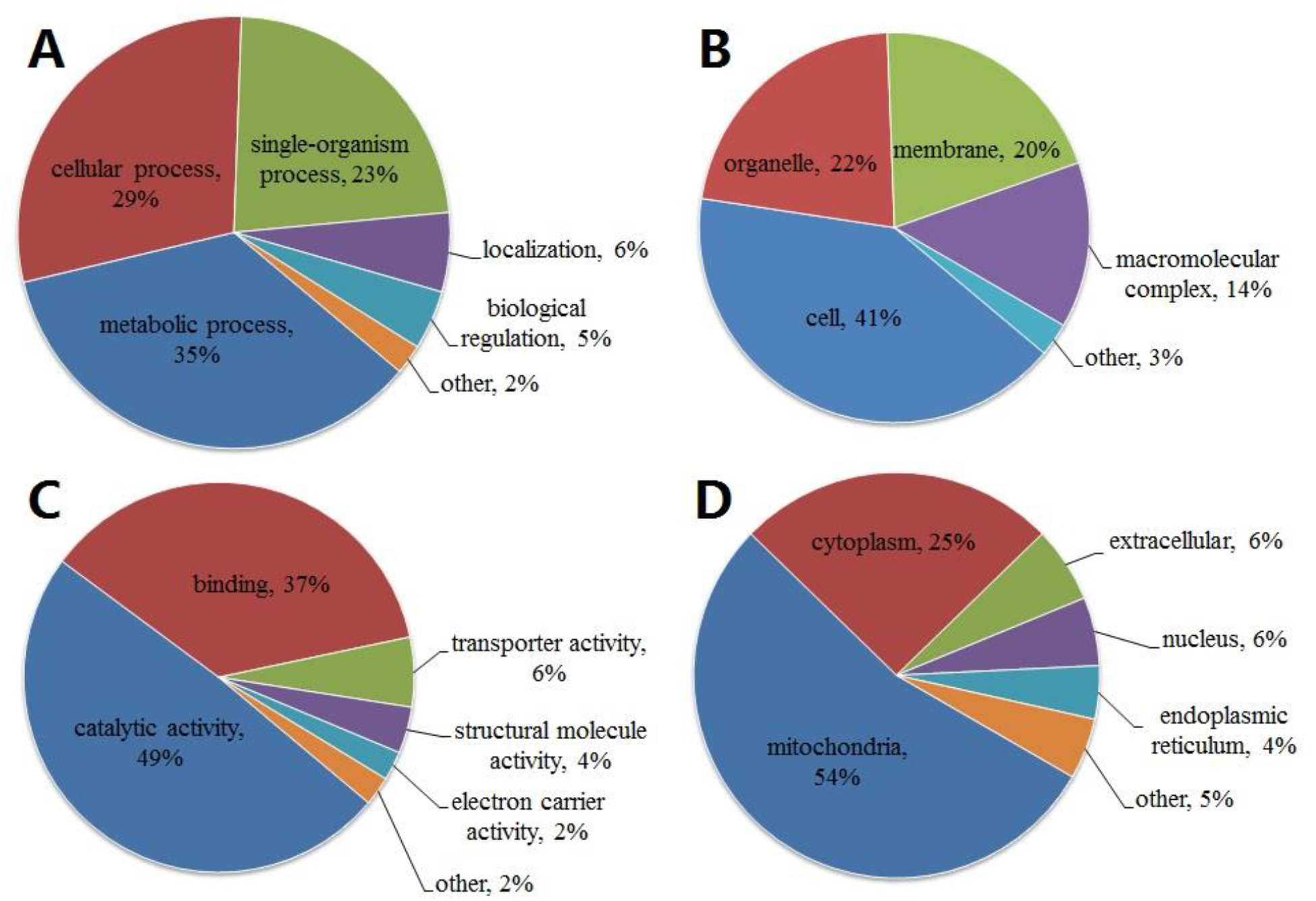
GO classification of the identified succinylated proteins in SRMV-infected vero cell A. Biological process; B. Cellular component; C. Molecular function; D. Subcellular localization

**Fig.7.**
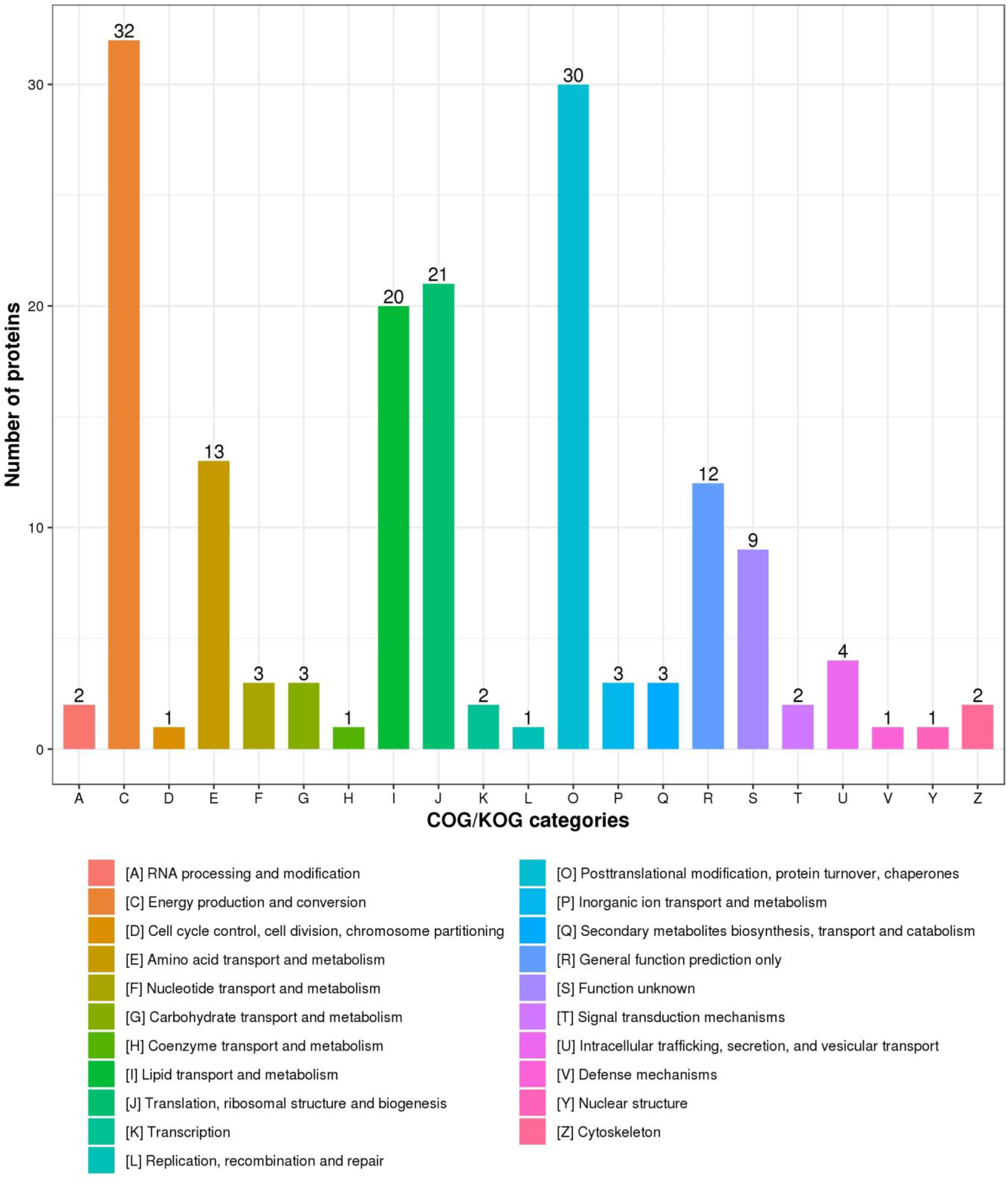
COG/KOG classification.

### Functional enrichment of the differential succiylated proteins

To further explore the functions of the succinylated proteins under SRMV infection, GO enrichment (cellular component, biological process and molecular function), KEGG and protein domain analysis were performed for all the identified succinylated proteins (Fig. 8, Supplementary Table S4).

**Fig.8.**
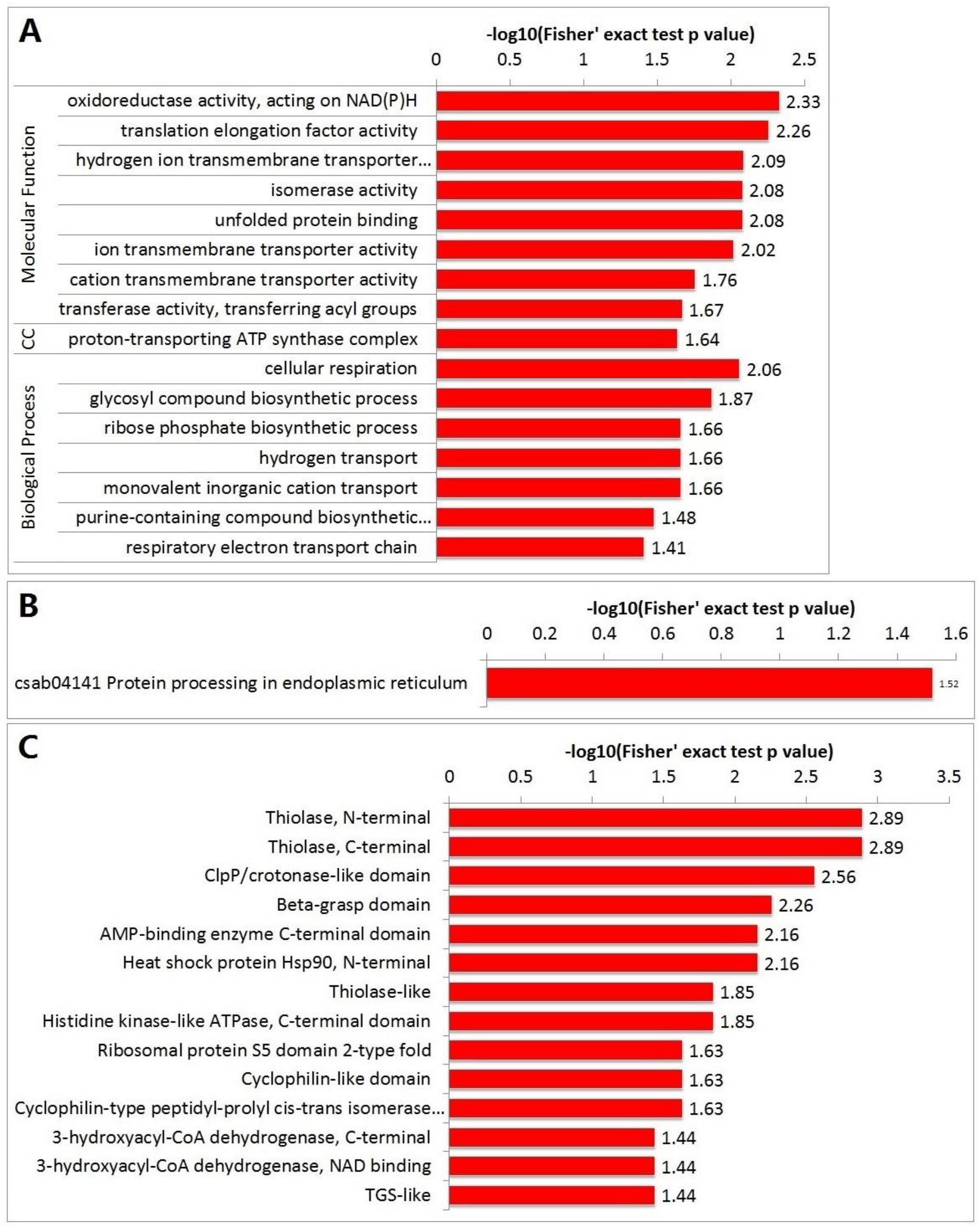
Enrichment of the succinylated proteins related to SRMV in vero cell. A. GO enrichment; B. KEGG pathway enrichment; C. Domain enrichment

The GO enrichment results of succinylated proteins were shown in Fig.8A. The cellular components were mainly enriched in proton-transporting ATP synthase complex. In the molecular function category, transmembrane transporter activity and unfolded protein binding were found to be significantly enriched. Lysine succinylated proteins were significantly enriched in the cellular metabolic process with specific enrichment in cellular respiration, ribose phosphate biosynthetic process and hydrogen transport. The KEGG database was used to identify the pathway involvement of these DScPs (Fig.8C). 13 of the DScPs were involved in protein processing in the endoplasmic reticulum (ER), and 9 of the 13 proteins were down-regulated. Furthermore, the protein domain analysis showed that a large proportion of succinylated proteins were enriched in ClpP/crotonase-like domain and ribosomal protein S5 (RPS5) domain 2-type fold (Fig. 8B).

### Protein–Protein interaction network analysis of the differentially succinylated proteins

PPIs are critical for various biological processes. To better understand how succinylation regulates diverse metabolic processes and cellular functions in SRMV-infected Vero cells, the PPI analysis was performed by searching the STRING database and PPI networks were visualized using Cytoscape software.

PPI network included 108 succinylated proteins as nodes, connected by 804 identified interactions (Fig. 9 and Supplementary Table S5). Two highly connected subnetworks, carbon metabolism and protein processing in the ER of DScPs were enriched. In the first subnetwork, 16 succinylated proteins with 133 Ksc sites and 215 direct physical interactions participated in carbon metabolism, suggesting that they could play key roles in energy transformation. Among the 16 succinylated proteins, four subunits of ATP synthases were identified as DsCPs in SRMV-infected cells: γ, β, O and δ. ATP synthases are molecular motors that enable conversion of energy. The second subnetwork comprised 6 succinylated proteins with 142 Ksc sites and 39 direct physical interactions and were related to protein disulfide-isomerase (PDI) and heat shock protein (Hsp), which are essential to stabilize the three-dimensional structure of proteins. In these two subnetworks, tight protein–protein interaction networks including 3 up-regulated proteins and 19 down-regulated proteins, were significantly enriched. Most of the proteins in the PPI network contain more than two succinylated sites.

**Fig.9.**
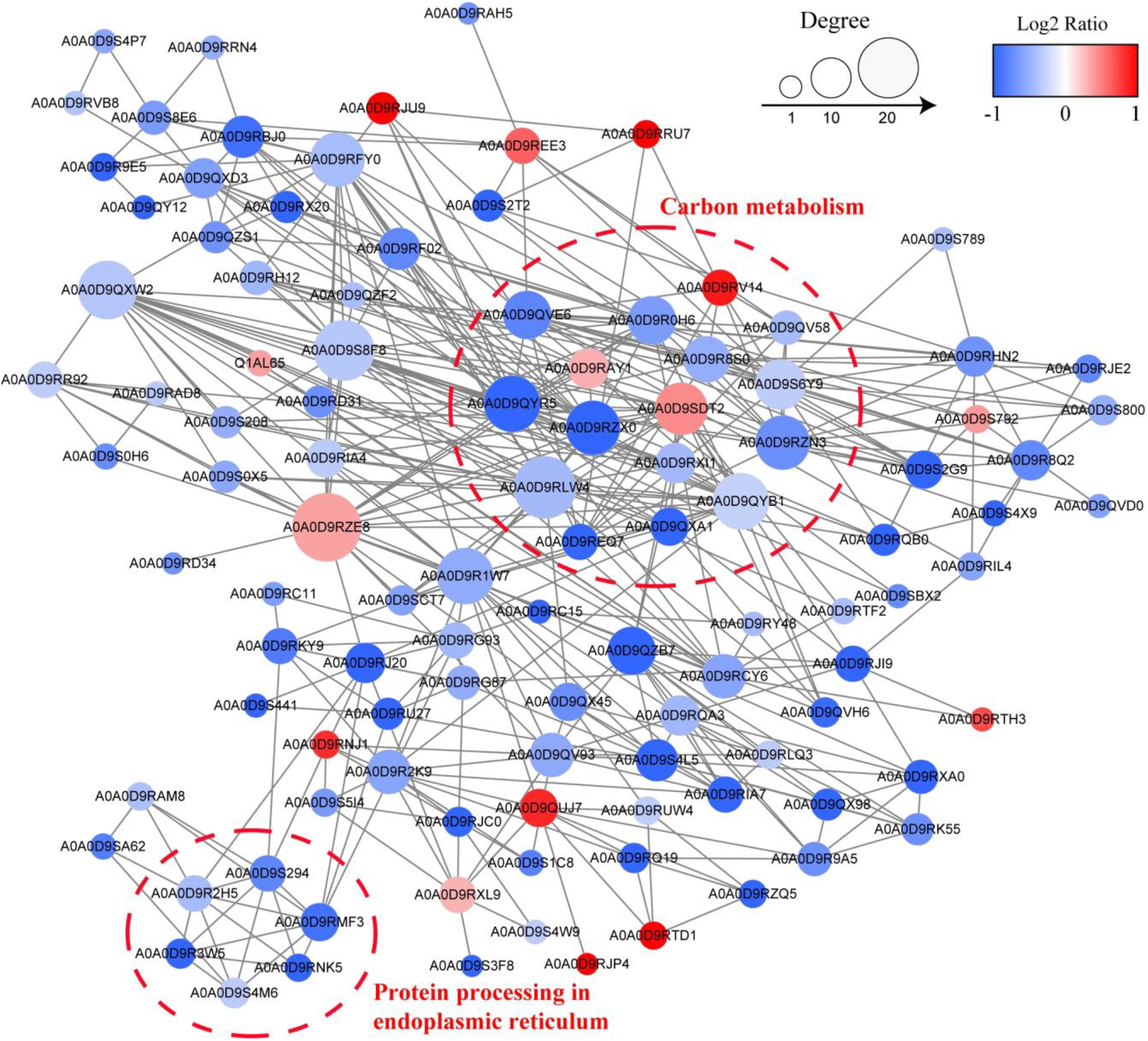
PPI networks of the succinylated proteins

To determine if the succinylation and acetylation occur at the same lysine site, we compared the succinylation data to the acetylation data (ProteomeXchange, PXD025081) of modification sites and proteins. The results showed that 34 succinylated proteins overlapped with acetylated proteins, one highly connected subnetwork, glyoxylate and dicarboxylate metabolism were enriched (Fig. 10). 18 succinylated sites of 13 proteins were also acetylated on the same lysine. Of these 13 proteins, 8 proteins had only one modification site, while 5 proteins had two modification sites. These results suggest that a complicated interaction among succinylated proteins might control the disease response or resistance during SRMV infection.

**Fig.10.**
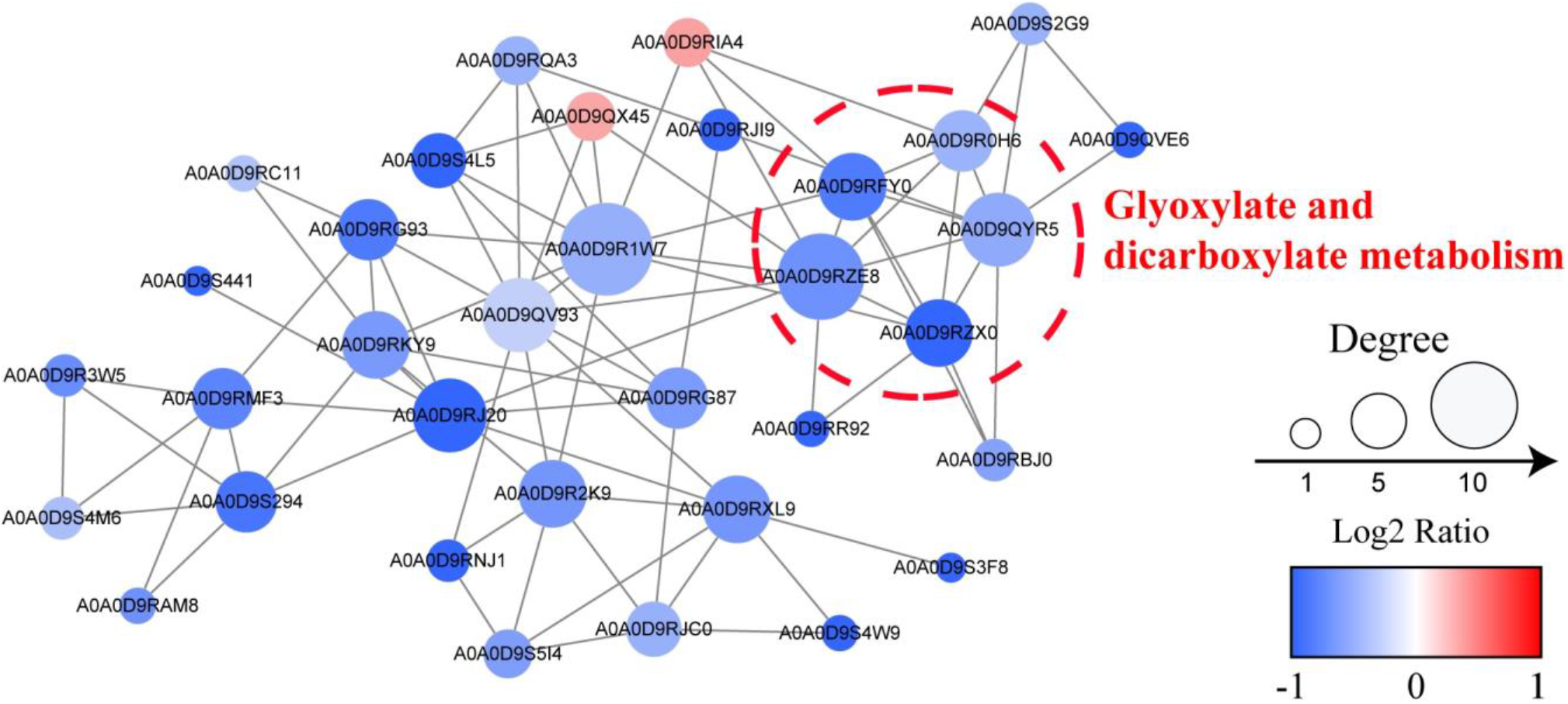
The crosstalk between succinylated and acetylated proteins

## DISCUSSION

Host infected with pathogen usually undergoes a series of physiological and biochemical changes at the cellular and molecular levels. It can lead to major changes during transcriptional, post-transcriptional, translational, and post-translational levels. As a conserved PTM, lysine succinylation is widespread in diverse species (14, 25–28), and hundreds of protein succinylation sites are present and being actively investigated. Succinylation has a negatively charged carboxylate group that results in more distinct chemical and structural changes in lysine residues. Lysine succinylation plays a central role in regulating multiple biological processes, especially for metabolism (36). The changes in metabolism alter succinylation, then succinylation provides feedback on metabolism and crosstalk with other proteins that are important to pathology. Succinylation was initially discovered in mitochondrial proteins involved in the TCA cycle, fatty acid metabolism, and amino acid degradation (37). It implies that succinylation may help to integrate the responses to the diverse metabolic challenges. Nonetheless, lysine succinylation modulates the activities of enzymes and pathways by modifying catalysis or cofactor binding sites. In addition to physiological processes, lysine succinylation is also associated with pathophysiological processes and diseases (38).

Up to date, although comparative transcriptome and proteomic profiling were analyzed in bone marrow-derived dendritic cells (BMDCs) or peripheral blood mononuclear cells (PBMCs) stimulated with SRMV (39–41), little is known about lysine succinylation in cells infected by SRMV. In the present study, for the first time, we elucidated the effects of SRMV infection on the protein lysine succinylation profiles of Vero cells by combining proteomics and succinylome approaches. This systematic analysis will provide an important baseline for functional studies on the response of succinylated proteins to SRMV.

Overall, we identified 2840 succinylation sites in 875 cellular proteins, and 2633 modification sites in 823 proteins regulated in SRMV-infected Vero cells. 177 proteins with 272 succinylation sites were differentially succinylated in response to SRMV. The numbers of succinylated sites and proteins were less than that of acetylation (ProteomeXchange, PXD025081). These results implied that acetylation may be more important than succinylation in SRMV infection. Of these DScPs, Hsps, ATP synthase subunits, PDIA4 and vimentin were modified at multiple succinylation sites. Succinylated sites at the 129 th of vimentin, 81 st and 113 th of HspA5, 56 th and 601 st of HspA8, 394 th and 600 th of HspA9, 112 nd and 496 th of HspD1 overlapped with acetylation sites. PDIA4 can significantly enhance the replication of influenza viruses (42). Similar to the result of acetylation, all succinylated sites of PDIA 4 were significantly down-regulated in this study, which suggests that SRMV replication might destroy the cellular homeostasis. Vimentin plays important roles in viral infection by interacting with viral proteins (43–53). These results suggested that the succinylated proteins participate in SRMV infection.

Although NLR family member X1 (NLRX1) only has one succinylated site in the study, it is virtually worth researching in immune system function of virus infection. NLRX1 can negatively regulate type-I interferon, attenuate pro-inflammatory NF-kB signaling and modulate autophagy, cell death, and proliferation. NLRX1 expression affects HIV infections, but seems to act controversially (54, 55). NLRX1 interacts with the influenza PB1-F2 protein to protect macrophages from apoptosis and downregulates IFN-β and IL-6 production (56). During the process of KSHV reactivation, NLRX1 plays a positive role in KSHV lytic replication by negatively regulating IFN-β response (57). NLRX1 promotes HCV infection by recruiting PCBP2 to inhibit MAVS via K48-linked polyubiquitination (58). NLRX1 interacts with Nsp9 of porcine reproductive and respiratory syndrome virus (PRRSV) to restrict viral replication (59).

The results of GO functional classification in the present study were similar to that of the acetylome of SRMV-infected cells (ProteomeXchange, PXD025081). The identified succinylated proteins are also related to catalytic activity and binding. Proteins related to proton-transporting ATP synthase complex were highly enriched and agree with the molecular function category and protein domain enrichment results. ClpP/crotonase plays an important role in protein quality and homeostasis in the cell by functioning mainly in the disaggregation, unfolding and degradation of native as well as misfolded proteins (60). RPS5 plays an essential role in protein translation and is related to cell differentiation and apoptosis (61, 62). RPS5 combines with the internal ribosome entry site (IRES) of hepatitis virus to provide the initial stages of translation initiation on the viral RNA (63). However, the molecular mechanism of RPS5 is still unclear. The subcellular localization of the DScPs was different from the differentially expressed lysine-acetylated proteins. DScPs were mainly located in mitochondria and cytoplasm. ER is critical for protein synthesis and maturation and resides on many molecular chaperones that assist protein folding and assembly. Virus infection can alter ER and activate the unfolded-protein response to facilitate viral replication (64, 65). The KEGG pathway enrichment analysis showed that 13 of the DScPs might play a major role in protein processing in the ER of SRMV-infected Vero cells. Therefore, the succinylated proteins potentially play vital roles in virus replication, and the relationship between SRMV infection and DScPs requires further investigations.

PPIs are critical for various biological processes. The present study provided the first global PPI network of succinylated proteins induced by SRMV infection. The subnetworks of carbon metabolism were enriched. ATP synthases are membrane protein complexes closely related to respiration in mitochondria. They are molecular motors and can synthesize ATP by continuous rotation, and maintain the energy needed for metabolism. In the second subnetwork, some members of PDI and Hsp were enriched. PDI can catalyze the formation, breakage and rearrangement of disulfide bonds, and promote protein folding, which is essential to stabilize the three-dimensional structure of proteins. The imbalance of PDI expression or enzyme activity is closely related to a series of diseases. Hsps play a variety of physiological functions by binding to protein molecules. Hsps can help amino acid chains fold into the correct three-dimensional structure, clean up the mis-folded proteins, escort proteins to find target molecules. Hsps also participate in innate immune response. These results suggest that the succinylation of these proteins play an important role in protein anabolism of SRMV-infected Vero cells. Moreover, 18 succinylation sites on 34 proteins overlapped with acetylation sites. Minimal overlap of succinylation and acetylation sites indicates differential regulation of succinylation and acetylation (37). However, others report succinylation occurs at a low level and extensively overlaps with acetylation in prokaryotes and eukaryotes (28, 66). Overall, these results provided a foundation for investigating the role of succinylation alone and its overlap with acetylation in response to SRMV infection and also raise the possibility of crosstalk between the two PTMs. A comprehensive study is required to determine whether the relationship between succinylation and acetylation in SRMV infection is independent or competitive, or if they complement each other to promote a regulatory role.

Histone modification is one typical way of epigenetic modifications. Histone succinylation is an enzymatic PTM as are other histone codes in the nucleus (67). It has important roles in nucleosome unwrapping rate and DNA accessibility (68). Recently, it has been identified that succinylation plays a crucial role in physiological and pathological processes (17, 28). However, to date rarely have studies reported the effect of histone succinylation on virus infection. Increasing evidence indicates that histone acylation modifications have important roles in virus infection. Yuan Y et al. first reported that histone succinylation might promote HBV replication (69). Interestingly, significantly expression of histone succinylated protein was not found in this study which indicates that histone succinylation might have little effect on SRMV infection.

## MATERIALS AND METHODS

### Cell, Virus and Infection

Vero cells were maintained in authors’ laboratory, and cultured in DMEM medium (Sigma Aldrich, St Louis, MO, USA) supplemented with 10% fetal bovine serum, 100 IU/ml penicillin and 100 μg/ml streptomycin at 37 °C in 5% CO_2_ incubator. The PPRV vaccine strain Nigeria 75/1 was cultured in our laboratory, which was propagated by Vero cells (70). Vero cells were infected with PPRV at MOI of 1 or mock-infected with phosphate-buffered saline (PBS, 0.01M, pH7.4) at 37°C for 1 h. The MOI was confirmed according to the viral titers of Vero cell line. After infection, the virus inoculum was removed, and the fresh medium was then added to wells and incubated. The infected cells were harvested at 24, 48, 72 h post infection (hpi) and analysed by western blotting and pan anti-succinyllysine antibody (PTM-419, PTM Biolabs, Hangzhou, China), and non-inoculated cells served as control group. Three biological replicates of independent (three parallel) experiments were performed.

### Protein Extraction

The harvested cell samples were washed twice with cold phosphate-buffered saline (PBS). Then, each sample was sonicated on ice in lysis buffer (8 M urea, 1% Protease Inhibitor Cocktail, 3 μM TSA, 50 mM NAM and 2 mM EDTA). The resulting supernatants were centrifuged with 12,000 rpm for 10 min at 4 °C to remove the cell debris. The protein concentration was determined with BCA kit according to the manufacturer’s instructions.

### Trypsin Digestion and Dimethylation Labeling

The protein solution was reduced with 5 mM dithiothreitol for 30 min at 56 °C and subsequently alkylated with 11 mM iodoacetamide for 15 min at room temperature in darkness. For tryptic digestion, the protein samples were diluted to urea concentration less than 2M by adding 100 mM NH_4_CO_3_. Finally, trypsin was added at 1:50 trypsin-to-protein mass ratio for the first digestion overnight and 1:100 trypsin-to-protein mass ratio for a second 4 h-digestion at 37 °C. Then, the peptide was desalted by Strata X C18 SPE column (Phenomenex) and vacuum-dried. Peptide samples were resuspended in 0.1 M TEAB and labeled in parallel in different tubes by adding CH_2_O or CD_2_O to the control and infected samples, respectively. The reactions were mixed and further treated with NaBH_3_CN (sodium cyanoborohydride) for 2 h at room temperature, then desalted adding formic acid and vacuum dried.

### Enrichment of Succinylated Peptides

Lysine-succinylated peptides were enriched using agarose-conjugated pan anti-succinyllysine antibody (PTM Biolabs, Hangzhou, China). In brief, dried tryptic peptides dissolved in NETN buffer (100 mM NaCl, 1 mM EDTA, 50 mM Tris-HCl, 0.5% NP-40, pH 8.0) were incubated with pre-washed pan anti-succinyllysine (PTM-104, PTM Biolabs, Hangzhou, China) conjugated agarose beads at 4°C overnight with gentle oscillation. Then the beads were washed 4 times with NETN buffer and 2 times with ddH_2_O. The bound peptides were eluted 3 times from the beads with 0.1% trifluoroacetic acid (TFA; Sigma-Aldrich, Saint Louis, USA). The eluted fractions were pooled together and vacuum-dried. For LC-MS/MS analysis. The resulting peptides were desalted with C18 ZipTips (Merck Millipore, Billerica, USA) according to the manufacturer’s instructions and dried by vacuum centrifugation.

### LC-MS/MS Analysis

The enriched peptides were dissolved in solvent A (0.1% Formic Acid (FA) in 2% acetonitrile (ACN)), directly loaded onto a home-made reversed-phase analytical column (1.9 μm beads, 120 A pore, 15-cm length, 75 μm i.d.). The gradient was comprised of a four-step linear: from 9% to 25% solvent B (0.1% FA in 90% ACN) for 24 min, from 25% to 40% for 10 min, from 40% to 80% for 3 min, and maintained at 80% for the last 3 min on an EASY-nLC 1000 UPLC (Ultra Performance Liquid Chromatography) system at a constant flow rate of 700 nL/min.

The peptides eluting from the analytical column were ionized and subjected to tandem mass spectrometry (MS/MS) in Q Exactive™ Plus (Thermo Scientific) coupled online to the UPLC using NanoSpray Ionization (NSI) source. The electrospray voltage applied was 2.0 kV. Intact peptides were detected at a resolution of 70,000 with a scan range of 350–1800 m/z for full MS scans in the Orbitrap. Peptides were then selected for MS/MS using an NCE setting of 28 and ion fragments were detected at a resolution of 17,500. A data-dependent acquisition (DDA) that alternated between one MS scan followed by 20 MS/MS scans was applied for the top 20 precursor ions, with 15 s dynamic exclusion. Automatic gain control (AGC) was used to prevent overfilling of the ion trap and set at 5E4, with a fixed first mass of 100 m/z.

### Database Searches

The protein succinylation sites identification and quantification were processed using MaxQuant search engine (v.1.5.2.8) against *Chlorocebus sabaeus* database (19,228 sequences), concatenated with reverse database and common contaminants. Trypsin/P was specified as cleavage enzyme allowing up to 4 missing cleavages. The mass tolerance for precursor ions was set to 20 ppm in the first analysis and 5 ppm in the full search, and the fragment mass tolerance was set to 0.02 Da. Carbamido-methylation of cysteine (Cys) was specified as fixed modification, and oxidation on methionine, succinylation on protein N-terminal and succinylation on lysine were specified as variable modifications. The false discovery rate (FDR) thresholds and the minimum score for modified peptide were set to <1% and >40, respectively. The minimum peptide length was set at 7. All other parameters in MaxQuant were set to default values.

### Bioinformatics analysis

Gene Ontology (GO) annotation was derived from the UniProt-GOA database (http://www.ebi.ac.uk/GOA). Proteins were classified based on three categories: biological process, cellular component and molecular function. Protein domain annotation was performed using the InterProScan (http://www.ebi.ac.uk/InterProScan/) based on protein sequence alignment method, and the InterPro domain database (http://www.ebi.ac.uk/interpro/) was used. The Kyoto Encyclopedia of Genes and Genomes (KEGG) database (http://www.genome.jp/kegg/) was used to annotate and map the pathways. GO, protein domain and KEGG pathway enrichment analysis were performed using the DAVID bioinformatics resources 6.8. Wolfpsort (https://wolfpsort.hgc.jp/), a subcellular localization predication soft was used to predict subcellular localization. Amino acid sequence motifs (within ± 10 residues of the succinylated sites) were analyzed by motif-X. Motif-based clustering analyses were also performed, and cluster membership was visualized using a heat map. Functional interaction network analysis was performed using the STRING database (v.11.0), with a high confidence threshold of 0.7, and visualized by Cytoscape 3.7.1.

All proteomics data associated with this manuscript have been uploaded to ProteomeXchange with the dataset identifier PXD025025.

## SUPPLEMENTAL MATERIAL

Supplemental material is available online only.

SUPPLEMENTAL TABLE S1.

SUPPLEMENTAL TABLE S2.

SUPPLEMENTAL TABLE S3.

SUPPLEMENTAL TABLE S4.

## ACKNOWLEDGEMENTS

This work was supported by the National Key Research and Development Program of China (2016YFD0500108). All authors would like to thank PTM Biolabs Inc. (Hangzhou, China).

